# Photosynthesis and antioxidant metabolism modulate the low-temperature resistance of seed germination in maize

**DOI:** 10.1101/2021.12.09.471969

**Authors:** Aiju Meng, Daxing Wen, Chunqing Zhang

**Affiliations:** State Key Laboratory of Crop Biology, Agronomy College, Shandong Agricultural University, Tai’an, Shandong Province 271018, P. R. China

**Keywords:** Seed germination, Seedling growth, Low-temperature stress, Maize, Transcriptome

## Abstract

Spring maize is usually subjected to low-temperature stress during seed germination, which retards seedling growth even if under a suitable temperature. However, the mechanism underlying maize seed germination under low-temperature stress modulating seedling growth after being transferred to normal temperature is still ambiguous. In this study, we used two maize inbred lines with different low-temperature resistance (SM and RM) to investigate the mechanism. The results showed that the SM line had higher lipid peroxidation and lower total antioxidant capacity and germination percentage than the RM line under low-temperature stress, which indicated that the SM line was more vulnerable to low-temperature stress. Further transcriptome analysis revealed that seed germination under low-temperature stress caused down-regulation of photosynthesis related gene ontology (GO) terms in two lines. Moreover, the SM line displayed down-regulation of ribosome and superoxide dismutase (SOD) related genes, whereas genes involved in SOD and vitamin B6 were up-regulated in the RM line. Kyoto Encyclopedia of Genes and Genomes (KEGG) enrichment analysis revealed that photosynthesis and antioxidant metabolism related pathways played important roles in seed germination in response to low-temperature stress, and the photosynthetic system displayed a higher damage degree in the SM line. Both qRT-PCR and physiological characteristics experiments showed similar results with transcriptome data. Taken together, we propose a model for maize seed germination in response to low-temperature stress.

**One sentence summary:** Damage degree of photosynthesis and total antioxidant capacity (especially SOD activity) determine diverse low-temperature resistance among maize inbred lines at the germination stage.

## Introduction

Maize (*Zea mays* L.) originated in tropical and subtropical areas and is naturally sensitive to low-temperature stress, especially during seed germination. Low-temperature limits the spread and production of maize all over the world (Zhang et al., 2020). As spring maize, seed germination and seedling growth at an early stage are usually subjected to low-temperature stress. Despite the increasing average temperature, further seedling growth also seems to be affected, which may be due to the inability of plants to respond quickly to favorable environmental changes (Sowiński et al., 2005). However, the mechanism underlying maize seed germination under low-temperature modulating seedling growth after being transferred to normal temperature is still ambiguous.

Numerous studies have shown that cold stress at the seedling stage affects photosynthesis by reducing the activity of photosystem II (PSII) (Savitch et al., 2011). Moreover, cold stress also affects energy collection and preservation at different points, and the reduction degree of photosynthetic activity is various among different genotypes (Ensminger et al., 2006). The chloroplast ultrastructure of seedlings developed under low-temperature stress is disordered and cannot be repaired after recovering to favorable conditions (Grzybowski et al., 2019). However, the mechanism of low-temperature stress at the germination stage affecting photosynthesis remains unknown.

Low-temperature stress can increase the accumulation of reactive oxygen species (ROS) (Li et al., 2019b), which can cause lipid peroxidation, DNA damage, protein denaturation, carbohydrate oxidation, pigment decomposition, and enzyme activity damage (Bose et al., 2014). Low-temperature tolerance is varied among different genotypes, which may be related to the antioxidant system. The main ROS scavenging enzymes in plants include superoxide dismutase (SOD), peroxidase (POD), catalase (CAT), glutathione peroxidase (GPX), and ascorbate peroxidase (APX). These enzymes provide efficient approaches for cells to detoxify O_2_^−^ and H_2_O_2_ together with antioxidants glutathione and ascorbic acid (Romero-Puertas et al., 2006). There is a close relationship between glutathione and the cold resistance of maize (Prasad, 1997). Reduced glutathione (GSH), a non-enzymatic antioxidant, is essential for maintaining the redox state of cells (Noctor et al., 2012).

Sugars are essential for plant growth and are involved in response to stress (Zheng et al., 2010). Sucrose is a common substance for energy storage and osmotic regulation in plant cells. Sucrose can also form a sugar layer around the cells, which has a higher membrane phase transition temperature to prevent cell dehydration (Zhang and Bartels 2018). The increase of sucrose content was the initial reaction of plants exposed to cold conditions (Nägele and Heyer, 2013).

Previous studies have reported the mechanism underlying seedling growth in response to low-temperature stress at the seedling stage, which is important to improve the low-temperature resistance of plants (Ma et al., 2015; Zeng et al., 2021). However, spring maize is more susceptible to low-temperature stress at the germination stage than the seedling stage. In this study, two maize inbred lines with different low-temperature resistance were used to investigate the effects of seed germination under low-temperature stress on seedling growth. The physiological experiments showed that the RM line had significantly higher total antioxidant capacity and sucrose content than the SM line under low-temperature stress, which may partially explain the different low-temperature resistance phenotypes. Subsequently, we further explore the differences in low-temperature resistance at the transcriptome level. Taken together, the results provide new insights into maize seed germination in response to low-temperature stress.

## Results

### The SM line is more vulnerable to low-temperature stress

To investigate the effects of maize seed germination under low-temperature stress on subsequent seedling growth under normal temperature, we firstly detected some traits of two maize inbred lines with different low-temperature resistance. In seed testing, maize seeds germinating at 25°C for 4 d (NT treatment) are usually used for evaluating seed germination energy. In this study, two lines had similar germination energy and seedling length (Fig. 1A). When the temperature decreased to 13°C (approximately half of the normal temperature), seed germination was remarkably slow down. The RM line showed more radicle emergence than the SM line at 13°C for 4 d (Fig. 1B). Given the same accumulated temperature, maize seeds firstly germinated at 13°C for 4 d and then transferred to 25°C for 2 d (LNT treatment). At this time, the RM line had notably higher seedlings than the SM line (Fig. 1C). If seeds germinated at 13°C for 7 d, seedling growth was significantly suppressed (Fig. 1D). Moreover, the germination percentage of the SM line decreased about 60% at 13°C for 7 d, while there was no significant difference in germination percentage between seeds germinated at 13°C for 7 d and seeds germinated at 25°C for 7 d in the RM line (Fig. 1E). Therefore, these results suggested that the SM line is more vulnerable to low-temperature stress. Subsequently, we detected the changes in lipid peroxidation, total antioxidant capacity (TAC), and sucrose content. Although lipid peroxidation levels of the two lines were significantly higher under LNT than those under NT, the RM line had a lower level of increased lipid peroxidation than the SM line (Fig. 1F). TAC decreased in the SM line but increased in the RM line under LNT treatment compared with NT treatment (Fig. 1G). Moreover, the RM line had notably higher sucrose content under LNT than that under NT (Fig. 1H). Taken together, seeds germinated under low-temperature stress increased TAC and sucrose content in the RM line, which might be related to low-temperature resistance.

**Figure 1.**
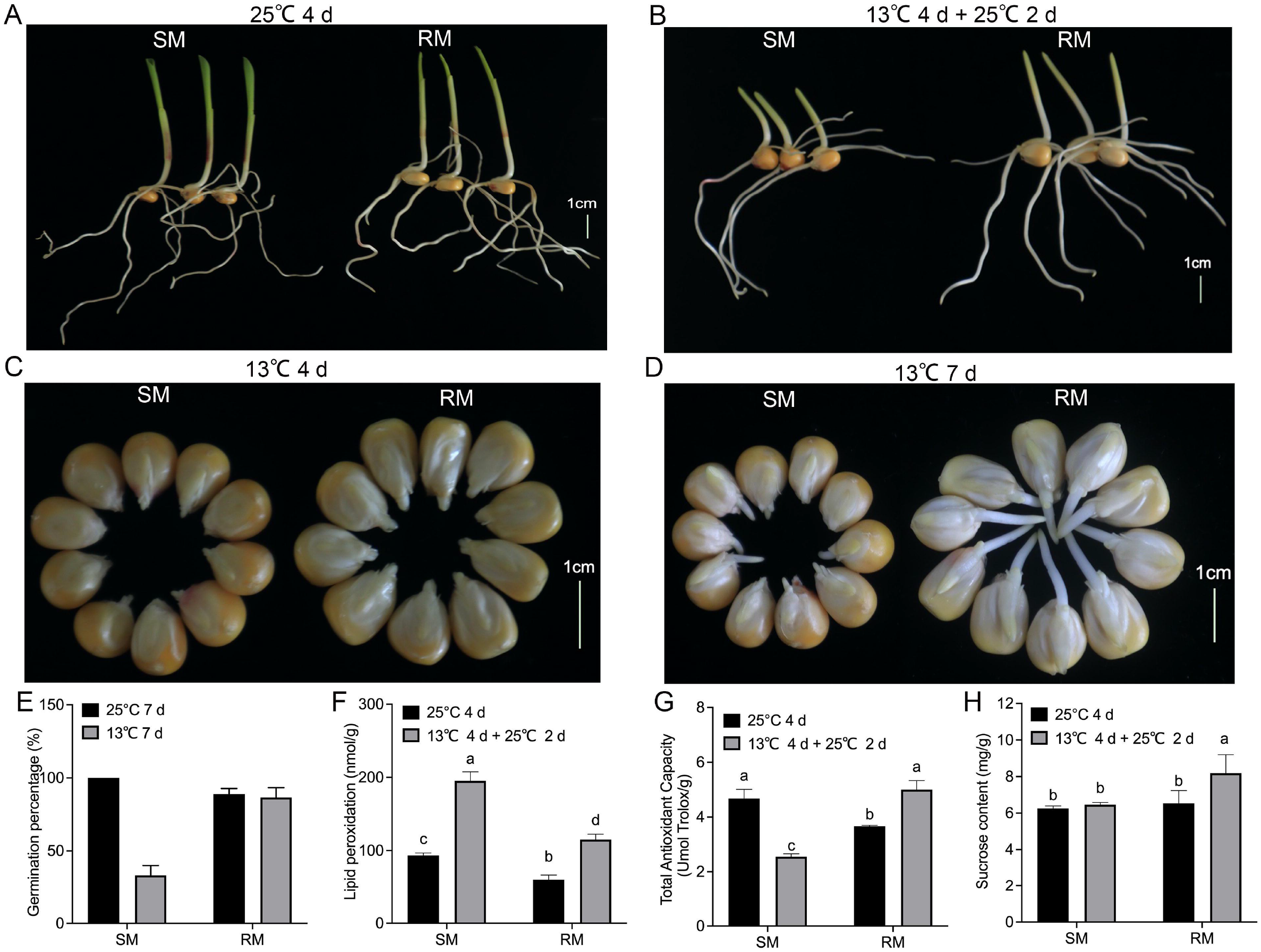
Effects of maize seed germination under low-temperature stress on subsequent seedling growth under normal temperature. A, Phenotypes of seed germination at 25°C for 4 d. B, Phenotypes of seed germination at 13°C for 4 d. C, Phenotypes of seed germination at 13°C for 4 d and then transferred to 25°C for 2 d. D, Phenotypes of 13°C at 13°C for 7 d. Scale bar, 1 cm. (E) Germination percentage. F, Lipid peroxidation. G, Total antioxidant capacity. (H) Sucrose content. Data are means ± SD (n = 3 replications of thirty plants). Different letters indicate significant differences among means under different treatments (p-value < 0.05).

### Transcriptome analysis of maize seed germination in response to low-temperature stress

To explore genes and metabolic pathways that control maize seed germination in response to low-temperature stress, we selected samples of RM and SM under NT and LNT for transcriptome analysis. Each treatment included three biological replicates. After removing adapters and sequences with low-quality regions, there remained approximately 39.83–50.39 million clean reads (Supplemental Table S1). Then, about 35.30-45.86 million clean reads were mapped to the maize genome. These clean reads included 83.34–88.45% uniquely mapped reads and 2.46–3.02% multiple mapped reads. DESeq2 R package was used to identify differentially expressed genes (DEGs) using padj < 0.05 and |log_2_Fold changeļ≥1 as the cutoff. The results displayed that 3186 genes were significantly upregulated and 4281 genes were significantly downregulated in the SM line under LNT compared with those under NT (SM_LNTvsNT), and 2797 genes were significantly upregulated and 3918 genes were significantly downregulated in the RM line under LNT compared with those under NT (RM_LNTvsNT) (Fig. 2A). Venn diagram showed that common up-regulated genes (54) were fewer than common down-regulated genes (608), most DEGs were specific in SM_LNTvsNT and RM_LNTvsNT (Fig. 2B and 2C).

**Figure 2.**
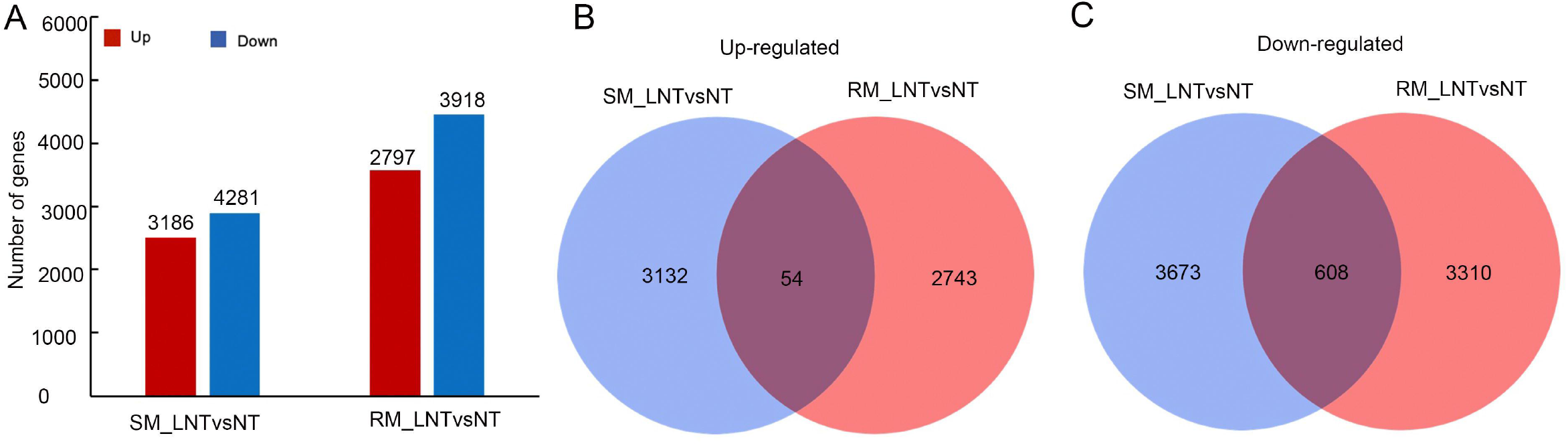
Differentially expressed genes in two maize inbred lines. A, The number of up-regulated and down-regulated DEGs between SM and RM lines. B, Venn diagram of up-regulated DEGs in SM_LNTvsNT and RM_LNTvsNT. C, Venn diagram of down-regulated DEGs in SM_LNTvsNT and RM_LNTvsNT. SM: low-temperature sensitive maize inbred line; RM: low-temperature resistant maize inbred line. NT treatment: seeds germinated at 25°C for 4 d. LNT treatment: maize seeds germinated at 13°C for 4 d and then transferred to 25°C for 2 d. SM_LNTvsNT: SM line samples under LNT compared with those under NT. RM_LNTvsNT: RM line samples under LNT compared with those under NT.

### Maize seed germination under low-temperature stress cause down-regulation of photosynthesis related GO terms

To gain insights into the functions of these DEGs, we first assigned gene ontology (GO) term(s) to each DEG with the GO database (version 2016.04, http://geneontol-ogy.org). Go enrichment analysis displayed that there was no significantly enriched GO term for common up-regulated genes, which might be due to the low number of DEGs. For these SM-specific up-regulated DEGs, there were only two significantly enriched GO terms, i.e. myosin complex (GO: 0016459, p = 4.82×10^−4^) in the cellular component group, ADP binding (GO: 0043531, p = 2.32×10^−5^) in the molecular function group (Fig. 3A). For these RM-specific up-regulated DEGs, there were nine significantly enriched GO terms. Among them, the most significantly enriched GO term was superoxide metabolic (GO: 0006801, p = 8.55×10^−6^) in the biological process group, vitamin binding (GO: 0019842, p = 5.35×10^−7^) in the molecular function group, which might play important roles in RM resistant to low-temperature stress (Fig. 3B).

**Figure 3.**
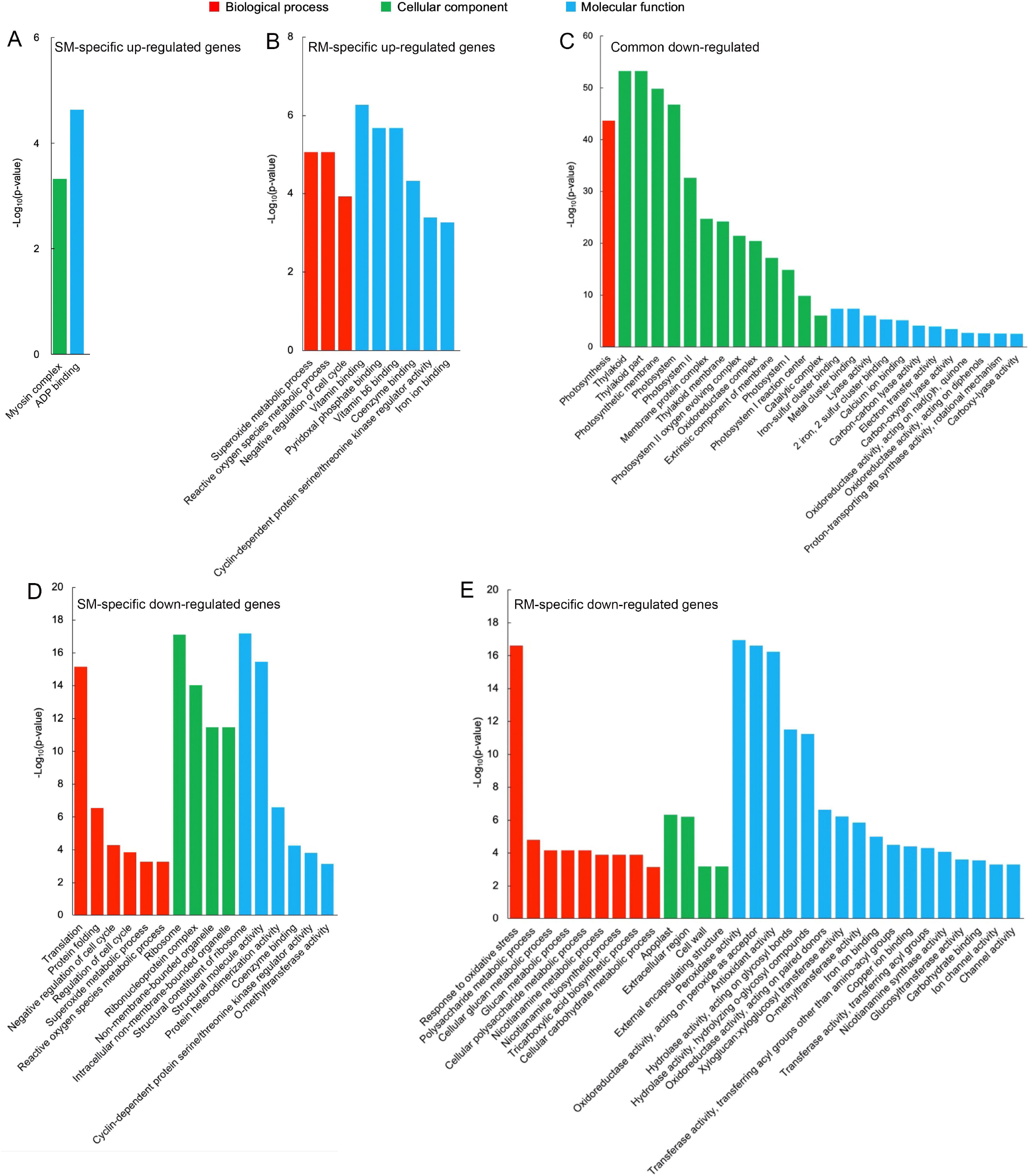
Significantly enriched GO terms in A, SM-specific up-regulated genes. B, RM-specific up-regulated genes. C, Common down-regulated genes. D, SM-specific down-regulated genes. E, RM-specific down-regulated genes in SM_LNTvsNT and RM_LNTvsNT. SM: low-temperature sensitive maize inbred line; RM: low-temperature resistant maize inbred line. NT treatment: seeds germinated at 25°C for 4 d. LNT treatment: maize seeds germinated at 13°C for 4 d and then transferred to 25°C for 2 d. SM_LNTvsNT: SM line samples under LNT compared with those under NT. RM_LNTvsNT: RM line samples under LNT compared with those under NT.

Compared with the up-regulated GO terms, there were more down-regulated GO terms. For these common down-regulated DEGs, the most significantly enriched GO term was photosynthesis (GO: 0015979, p = 2.16×10^−44^) in the biological process group, thylakoid (GO: 0009579, p = 5.33×10^−54^) in the cellular component group, iron-sulfur cluster binding (GO:0051536, p = 4.38×10^−8^) in the molecular function group (Fig. 3C). Moreover, many photosynthesis related GO terms were also enriched. The results indicated that maize seed germination under low-temperature stress caused downregulation of photosynthesis related GO terms in both SM and RM lines. Translation (GO: 0006412, p =6.79×10^−16^) in the biological process group, ribosome (GO: 0005840, p = 7.64×10^−18^) in the cellular component group, structural constituent of ribosome (GO: 0003735, p = 6.47×10^−18^) in the molecular function group, represented the most markedly enriched GO terms in SM-specific down-regulated DEGs, suggesting that ribosomes of SM line might be damaged by low-temperature stress (Fig. 3D). For RM-specific down-regulated DEGs, the most prominently enriched GO terms were response to oxidative stress (GO: 0006979, p = 2.41×10^−17^) in the biological process group, apoplast (GO: 0048046, p = 4.58×10^−7^) in the cellular component group, peroxidase activity (GO: 0004601, p = 1.14×10^−17^) in the molecular function group (Fig. 3E). The results indicated that peroxidase activity might be not involved in the enhanced antioxidant capacity of the RM line in response to low-temperature stress.

### Photosynthesis and antioxidant metabolism pathways are involved in response to low-temperature stress at the germination stage

To identify the metabolic or signal pathways involved in maize seed germination under low-temperature stress, we further analyzed the Kyoto Encyclopedia of Genes and Genomes (KEGG) enrichment pathways. The results showed that photosynthesis-antenna proteins, photosynthesis, phenylpropanoid biosynthesis, flavonoid biosynthesis, glutathione metabolism, porphyrin and chlorophyll metabolism, stilbenoid, diarylheptanoid, and gingerol biosynthesis were markedly enriched KEGG pathways for common DEGs (Fig. 4). Therefore, photosynthesis and antioxidant metabolism pathways might play important roles in response to low-temperature stress at the germination stage.

**Figure 4.**
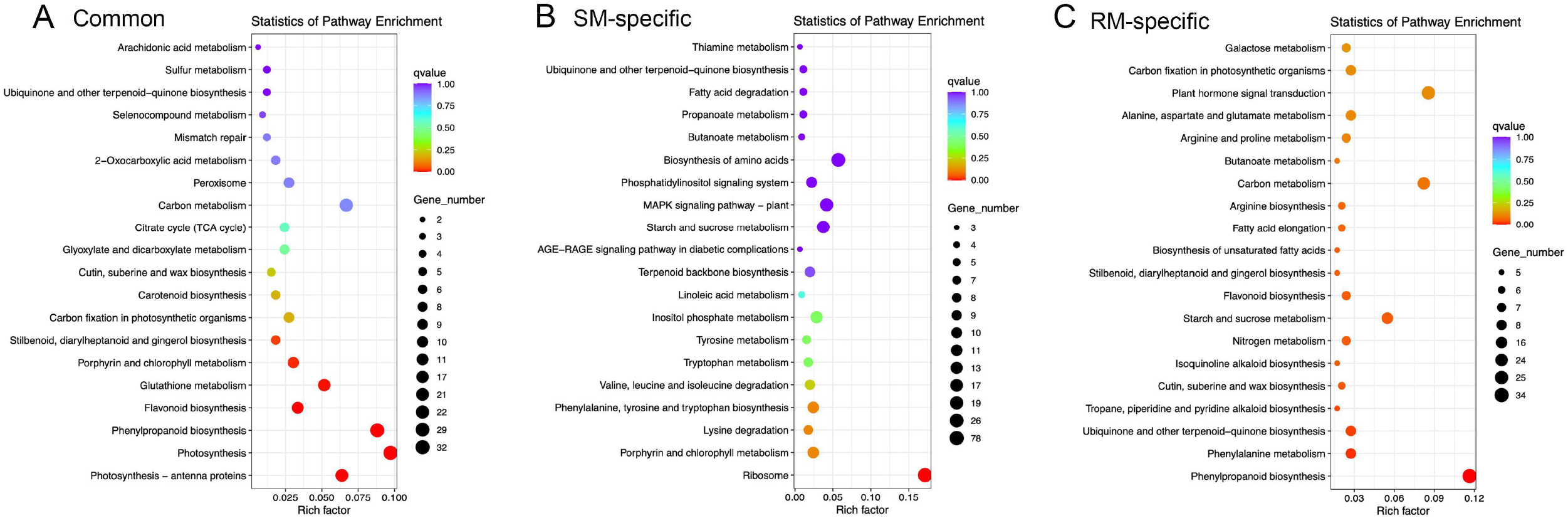
Top 20 KEGG pathways in A, Common DEGs. B, SM-specific DEGs. C, RM-specific DEGs in SM_LNTvsNT, and RM_LNTvsNT. SM: low-temperature sensitive maize inbred line; RM: low-temperature resistant maize inbred line. NT treatment:seeds germinated at 25°C for 4 d. LNT treatment: maize seeds germinated at 13°C for 4 d and then transferred to 25°C for 2 d. SM_LNTvsNT: SM line samples under LNT compared with those under NT. RM_LNTvsNT: RM line samples under LNT compared with those under NT.

Only the ribosome pathway was significantly enriched for SM-specific DEGs, which was also enriched in GO enrichment analysis. In the ribosome pathway, there were 78 DEGs in SM_LNTvsNT (Supplemental Table S2). Of the 78 DEGs, most genes were down-regulated except for three genes (*Zm00001d022111*, *Zm00001d022197*, and *Zm00001d002462*), which indicated that low-temperature stress might have a great influence on the ribosomal pathway of low-temperature sensitive inbred lines. For RM-specific DEGs, the significantly enriched KEGG pathways were related to phenylpropanoid biosynthesis, phenylalanine metabolism, ubiquinone, and other terpenoid-quinone biosynthesis, tropane, piperidine and pyridine alkaloid biosynthesis (Supplemental Table S3). In tropane, piperidine and pyridine alkaloid biosynthesis, only one gene was down-regulated in five DEGs.

### The photosynthetic system of the SM line is more vulnerable to low-temperature stress

The DEGs enriched in photosynthesis-related pathways were mainly located in the chloroplast and annotated to function as antenna proteins, photosystems I and II components, porphyrin, and chlorophyll metabolism related proteins (Fig. 5, Supplemental Table S4). Although DEGs involved in photosynthetic-antenna proteins were all down-regulated in both SM and RM lines, the down-regulation degree of the DEGs was lower in the RM line than the SM line. Moreover, most of the DEGs involved in photosynthesis and porphyrin and chlorophyll metabolism pathway were also down-regulated in both SM and RM lines, and the RM line also had a lower down-regulation degree of the DEGs than the SM line (Fig. 5, Supplemental Table S4). Taken together, the photosynthetic system of the SM line was even more badly damaged when seed germinated under low-temperature stress.

**Figure 5.**
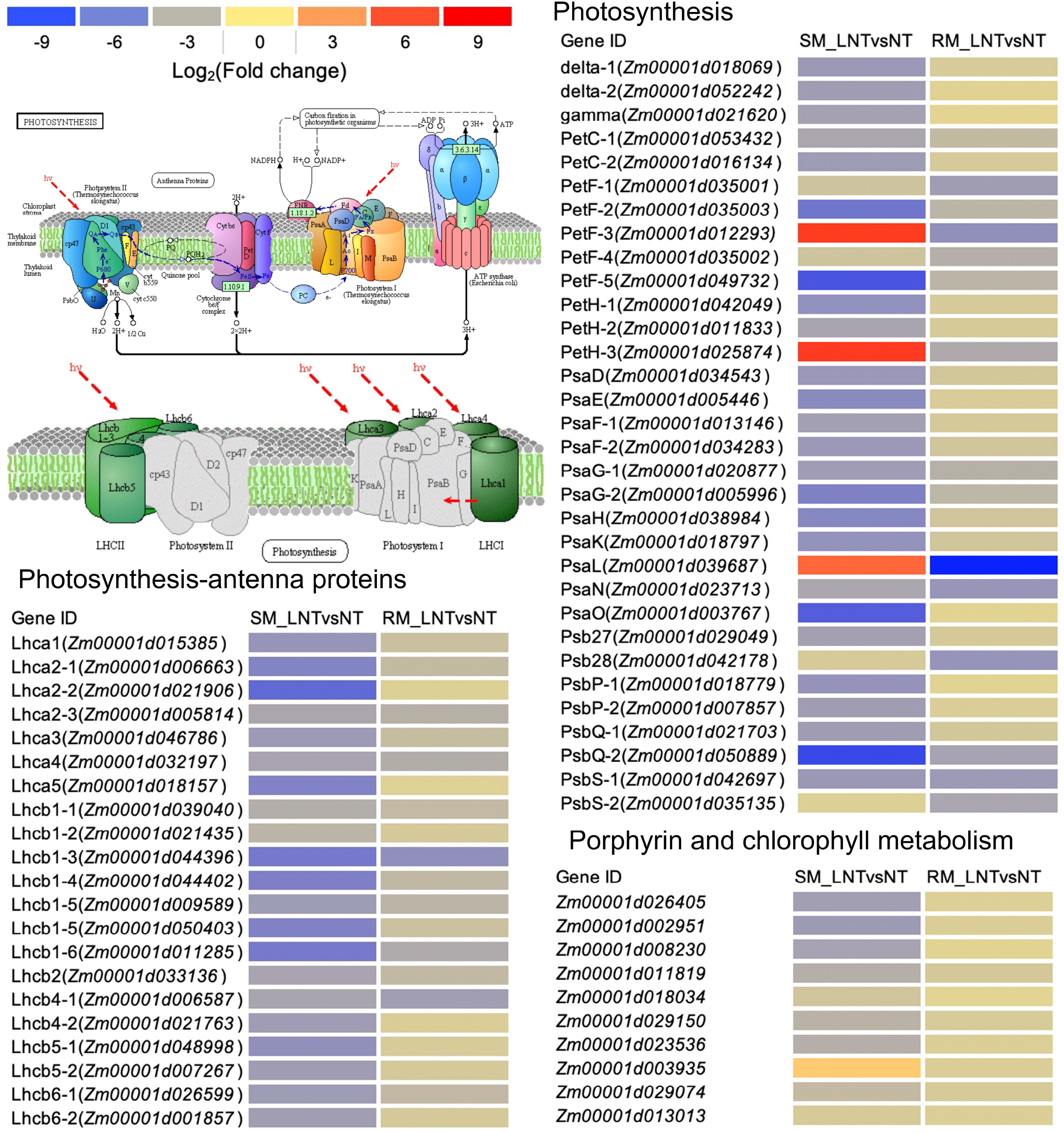
Heat map of photosynthesis related genes enriched in KEGG pathways in SM_LNTvsNT and RM_LNTvsNT. SM: low-temperature sensitive maize inbred line; RM: low-temperature resistant maize inbred line. NT treatment: seeds germinated at 25°C for 4 d. LNT treatment: maize seeds germinated at 13°C for 4 d and then transferred to 25°C for 2 d. SM_LNTvsNT: SM line samples under LNT compared with those under NT. RM_LNTvsNT: RM line samples under LNT compared with those under NT. Detailed lists of the DEGs are shown in Additional file Table S3. The color code from blue to red suggests the expression level of the DEGs normalized as the log_2_(Fold change).

### Validation of transcriptome data by qRT-PCR and physiological characteristics

To validate the gene expression level of the DEGs identified by RNA-seq, we randomly selected six DEGs to perform qRT-PCR assays (Fig. 6). All the DEGs showed similar expression patterns in the qRT-PCR assays as their changes of relative expression level identified by RNA-seq, suggesting the transcriptome data were credible. By extending the seedling growth at 25□ for 2 d, we observed that maize seed germination under low-temperature markedly inhibited subsequent seedling growth under normal temperature, especially in the SM line (Fig. 7A and 7B). Subsequently, we further detected the changes in SPAD value, SOD, and POD activities. Both SM and RM lines displayed a significant decrease of SPAD value under LNT and the decrease degree in the SM line was larger than that in the RM line, which was consistent with the changes of the photosynthetic system from transcriptome analysis (Fig. 7C). The SM line had lower SOD activity and higher POD activity under LNT than NT treatment, while the RM line showed the opposite trend of SOD and POD activities, which were consistent with the results of GO enrichment analysis (Fig. 7D and 7E). Moreover, the changes in SOD activities in both SM and RM lines were similar to the trends of TAC (Fig. 1F and 7D). Therefore, SOD activity might play a key role in TAC in seed germination under low-temperature stress.

**Figure 6.**
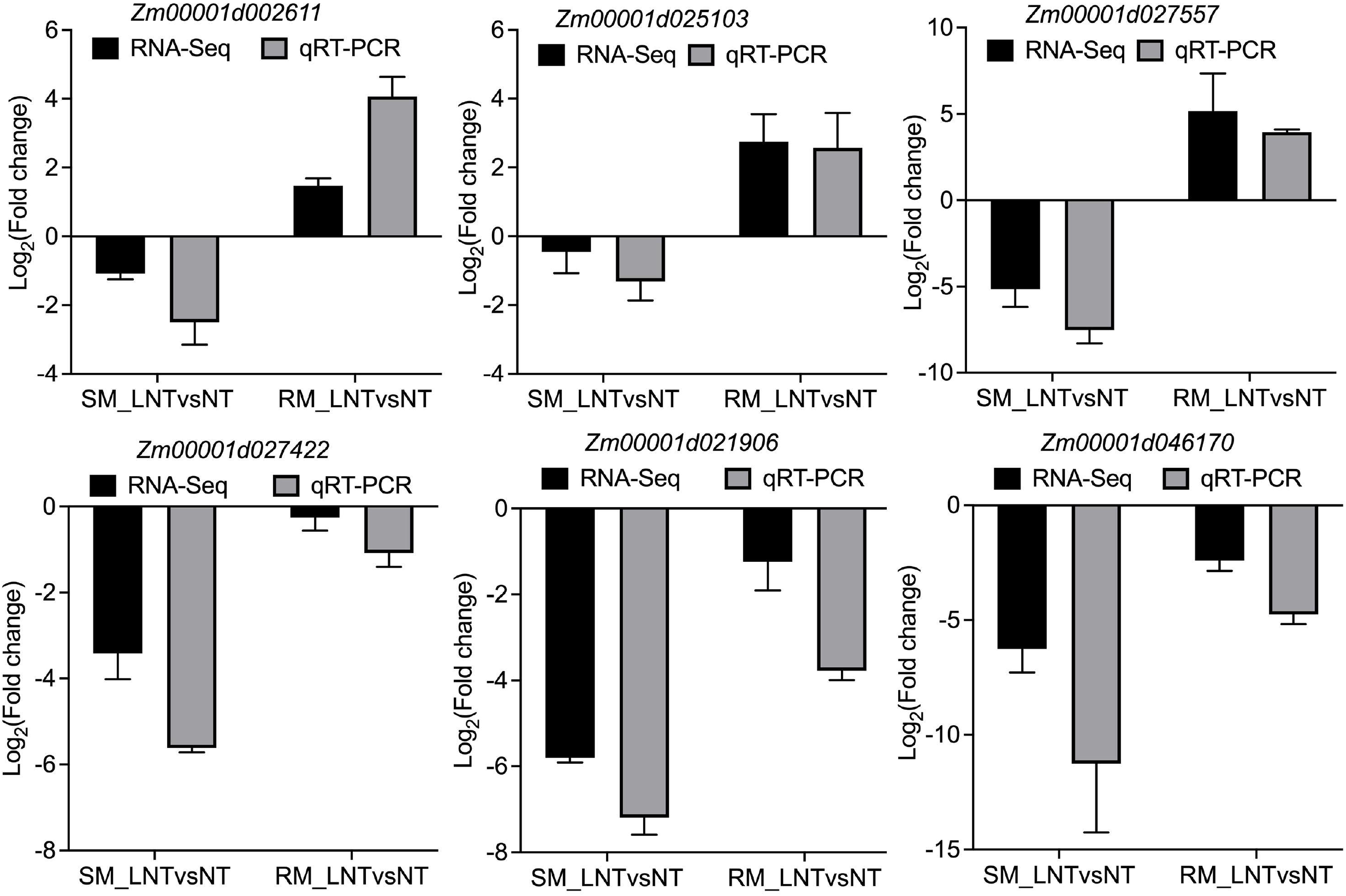
Validation of DEGs by qRT-PCR. The black bar and grey bar represent the fold change of the relative expression level from the RNA-seq data and qRT-PCR data, respectively. The maize *Actin* gene, as an internal control, was used to normalize the expression levels of the target genes. The error bars represent the standard deviations of three replicates. SM: low-temperature sensitive maize inbred line; RM: low-temperature resistant maize inbred line. NT treatment: seeds germinated at 25°C for 4 d. LNT treatment: maize seeds germinated at 13°C for 4 d and then transferred to 25°C for 2 d. SM_LNTvsNT: SM line samples under LNT compared with those under NT. RM_LNTvsNT: RM line samples under LNT compared with those under NT.

**Figure 7.**
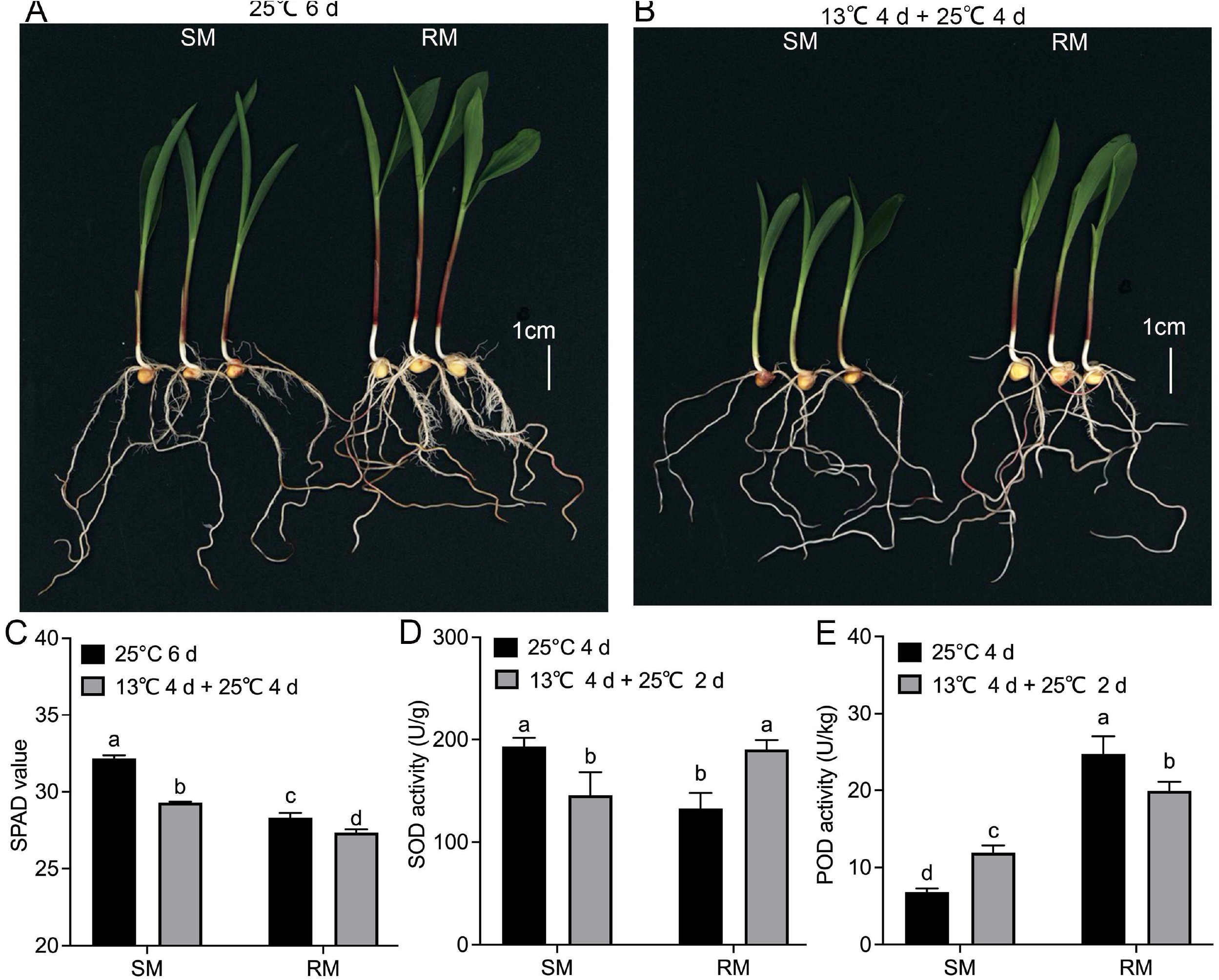
Validation of transcriptome data by physiological characteristics. A, Phenotypes of seed germination at 25°C for 6 d. B, Phenotypes of seed germination at 13°C for 4 d and then transferred to 25°C for 4 d. C, SPAD value. D, Superoxide dismutase. E, Peroxidase. Data are means ±SD (n = 3 replications of thirty plants). Different letters indicate significant differences among means under different treatments (p-value < 0.05). SM: low-temperature sensitive maize inbred line; RM: low-temperature resistant maize inbred line.

## Discussion

Low-temperature stress often occurs at the germination stage, which retards the seedling emergence of spring maize. Maize is vulnerable to low-temperature stress at the germination stage and the early stage of seedling establishment (Zhang et al., 2020). Deciphering the mechanism underlying maize seed germination in response to low-temperature stress can help for improving low-temperature resistance.

ROS generation in plant cells can be induced by some environmental factors, such as cold (Li et al., 2019b), drought (Zheng et al., 2020), heat (Zhao et al., 2018), and cadmium stress (Gu et al., 2019). ROS, as signal molecules, trigger signal transduction pathways in response to these abiotic stresses. In addition, ROS can cause irreversible cell damage through its strong oxidation characteristics, thereby promoting the change of plant morphology and structure, and enhancing resistance (Ohama et al., 2017; Bose et al., 2014). ROS can cause lipid peroxidation, DNA damage, protein denaturation, carbohydrate oxidation, pigment decomposition, and enzyme activity damage (Bose et al., 2014). In the present study, both SM and RM lines showed enhanced lipid peroxidation in seed germination under low-temperature stress, which were consistent with previous studies.

The detoxification mechanism of ROS plays an important role in the normal metabolism of plants, especially under stress. The main ROS scavenging enzymes in plants include SOD, POD, CAT, GPX, and APX. Previous studies have shown that the activities of CAT and monodehydroascorbate reductase are effective screening tools for maize hybrids with cold resistance (Hodges et al., 1997). The activities of antioxidant enzymes significantly increase when maize seeds germinate under low-temperature (Cao et al., 2019). In the present study, the RM line under LNT displayed increased SOD activity and decreased POD activity compared with NT treatment, while the SM line showed opposite trends in both SOD and POD activities (Fig. 7D-F). Moreover, GO enrichment analysis showed similar trends with the activities of SOD and POD (Fig. 3B and 3E). Interestingly, the changes in TAC were consistent with SOD activity (Fig. 1F and 7D). Therefore, SOD activity might play a key role in the TAC of maize seed germination under low-temperature stress. A previous study has shown that the intrinsic high level of SOD in halophytes is necessary to trigger a series of adaptive responses, and the role of other enzymatic antioxidants might reduce the basic level of H_2_O_2_ (Bose et al., 2014). Whether SOD activity also triggers a series of adaptive responses under low-temperature stress needs further study.

Glutathione metabolism is involved in regulating redox-sensitive signal transduction of plant tissues and maintaining antioxidant properties (Cnubben et al., 2001; Noctor et al., 2012). Compared with NT treatment, glutathione metabolism-related genes encoding glutathione peroxidase (*Zm00001d026154* and *Zm00001d002704*), glutathione transferase (*Zm00001d018220*, *Zm00001d027557*, *Zm00001d042102*, *Zm00001d029706* and *Zm00001d043344*), isocitrate dehydrogenase (*Zm00001d044021*) were all up-regulated only in RM line under LNT (Supplemental Table S5). Of these genes, *Zm00001d026154* and *Zm00001d029706* have been reported that they can help maize to resist drought stress (Zheng et al., 2020). Therefore, *Zm00001d026154* and *Zm00001d029706* might be essential for maize resistance to various abiotic stresses.

Vitamin B6 contains six forms, pyridoxal, pyridoxamine, pyridoxine, pyridoxal 5’-phosphate (PLP), pyridoxamine 5’-phosphate, and pyridoxine 5’-phosphate, of which PLP is the active form (Denslow et al., 2007). PLP is essential for many biochemical reactions, including decarboxylation, transamination, deamination, racemization, and trans sulfur reactions, which are mainly related to amino acid synthesis (Drewke and Leistner, 2001). Vitamin B6 as a cofactor has been fully confirmed. Moreover, vitamin B6 as an effective antioxidant and a factor that can increase resistance to biotic and abiotic stress has been proved (Huang et al., 2013). Previous studies have shown that VB6 is an effective singlet oxygen quencher, and its quenching rate is equivalent to or higher than that of vitamin C and E which are known as the two most effective biological antioxidants (Ehrenshaft et al., 1999; Ehrenshaft et al., 1998; Denslow et al., 2007). In the present study, GO enrichment analysis of RM-specific up-regulated DEGs showed that pyridoxal phosphate binding (GO: 0030170, p = 2.08×10^−6^) and vitamin B6 binding (GO: 0070279, p = 3.85×10^−8^) were significantly enriched in the molecular function group (Fig. 3B). Therefore, vitamin B6 might be involved in enhancing the low-temperature resistance of maize at the germination stage.

Photoinhibition occurs when the harvested light energy exceeds the available energy of chloroplasts or low-temperature sensitive plants exposed to low-temperature stress (Li et al., 2019b). Moreover, low-temperature stress can regulate PSII activity, leading to the loss of photosynthetic capacity (Savitch et al., 2011). The down-regulation of light-harveting complex protein will affect the downstream energy-related processes, and ultimately affect the overall growth and development of plants (Savitch et al., 2011). In this study, most genes related to photosynthetic apparatus were down-regulated in both RM and SM lines, but the decrease degree of the SM line was greater than that of the RM line (Fig. 5). Compared with NT treatment, the RM line shows better recovery ability in photosynthesis than the SM line under LNT. Cold affects photosynthesis through overexcitation of PSII reaction centers and the production of oxygen free radicals (Nie et al., 1992). ROS has harmful effects on photosynthetic devices (Fenza et al., 2017). In this study, the RM line has higher TAC than the SM line, which might be related to the smaller decrease in photosynthesis in the RM line. After low-temperature treatment, the SPAD values of the two lines decreased significantly, but the decrease degree of the SM line was greater than that of the RM line, which was consistent with the transcriptome data. Therefore, seed germination under low-temperature stress reduced subsequent seedling photosynthesis, but the decreased degree of photosynthesis was different between maize inbred lines with various low-temperature resistance.

Ribosomes are implicated in resistance to various adverse conditions (Garcia-Molina et al., 2020). A large number of down-regulated DEGs were enriched in the ribosome (GO: 0005840, p = 7.64×10^−18^) in the SM line (Supplemental Table S2). In mammalian cells, nucleolus, especially ribosome, is considered to be the hub of integrating cell response to adverse conditions(Yang et al., 2018; Pfister 2019). Some plant species have similar regulatory roles of ribosomes under abiotic and biotic stresses (Garcia-Molina et al., 2020). So far, a large number of ribosomal proteins (RPS) mutants with defects in chloroplast ribosomes have been reported in plants (Schultes et al., 2000; Xu et al., 2013; Zhang et al., 2016). In the present study, gene (*Zm00001d012353*) encoding 30S ribosomal protein S17 chloroplastic was significantly down-regulated in SM_LNTvsNT. The first plant plastid ribosomal protein mutant (high chlorophyll fluorescence 60) in maize displays an unstable light green effect on seedling growth due to the lack of plastid ribosomal small subunit protein 17 (Schultes et al., 2000). The transcription level of RPS is up-regulated after low-temperature acclimation (Garcia-Molina et al., 2020). Why is RPS transcription up-regulated after cold acclimation? The up-regulation of RPS under low-temperature stress is considered to maintain the rate of protein synthesis under thermodynamic adverse conditions (Garcia-Molina et al., 2020). In this study, genes encoding 30S ribosomal protein S1 chloroplastic (*Zm00001d038835*), 30S ribosomal protein S10 chloroplastic (*Zm00001d028153*), 30S ribosomal protein S4 chloroplastic (*Zm00001d047186*), 30S ribosomal protein S6 alpha chloroplastic (*Zm00001d034808*), 50S ribosomal protein L1 chloroplastic (*Zm00001d038084*), 50S ribosomal protein L11 chloroplastic (*Zm00001d027421*), 50S ribosomal protein L17 chloroplastic (*Zm00001d012998*), 50S ribosomal protein L21 chloroplastic (*Zm00001d053377*), 50S ribosomal protein L6 chloroplastic (*Zm00001d047462*) were all down-regulated only in the SM line. The role of translation and ribosome in adaptation to abiotic environment changes has been found in a recent analysis of the corresponding Arabidopsis mutants (Reiter et al., 2020). In particular, various examples of impaired cold tolerance due to the inactivation of chloroplast proteins involved in translation have been described, including subunits (Wang et al., 2017), biogenesis factors (Reiter et al., 2020) of the plastid ribosome-associated proteins (Pulido et al., 2018), translation initiation or elongation factors (Liu et al., 2010) and RNA-binding proteins (Kupsch et al., 2012). Therefore, down-regulated chloroplastic ribosomal protein-related genes in the SM line under LNT might cause the down-regulation of genes involved in the photosystem, thereby affecting photosynthesis and decreasing SPAD value.

Overall, we propose a possible network of maize seed germination under low-temperature stress affecting subsequent seedling growth (Fig. 8). Maize seed germination under low-temperature stress caused down-regulation of photosynthesis related genes in both SM and RM lines, but the SM line showed a larger decrease in the photosynthetic system than the RM line. Moreover, the SM line displayed down-regulation of the ribosome and SOD related genes, whereas genes involved in SOD and vitamin B6 were up-regulated in the RM line. SOD activity might play a key role in the TAC of maize seed germination under low-temperature stress because that the changes in TAC were consistent with SOD activity. The inhibition of maize seed germination under low-temperature on seedling growth might be mainly due to impaired photosynthesis. The differences of TAC (especially SOD activity) among various lines might affect low-temperature resistance at the germination stage.

**Figure 8.**
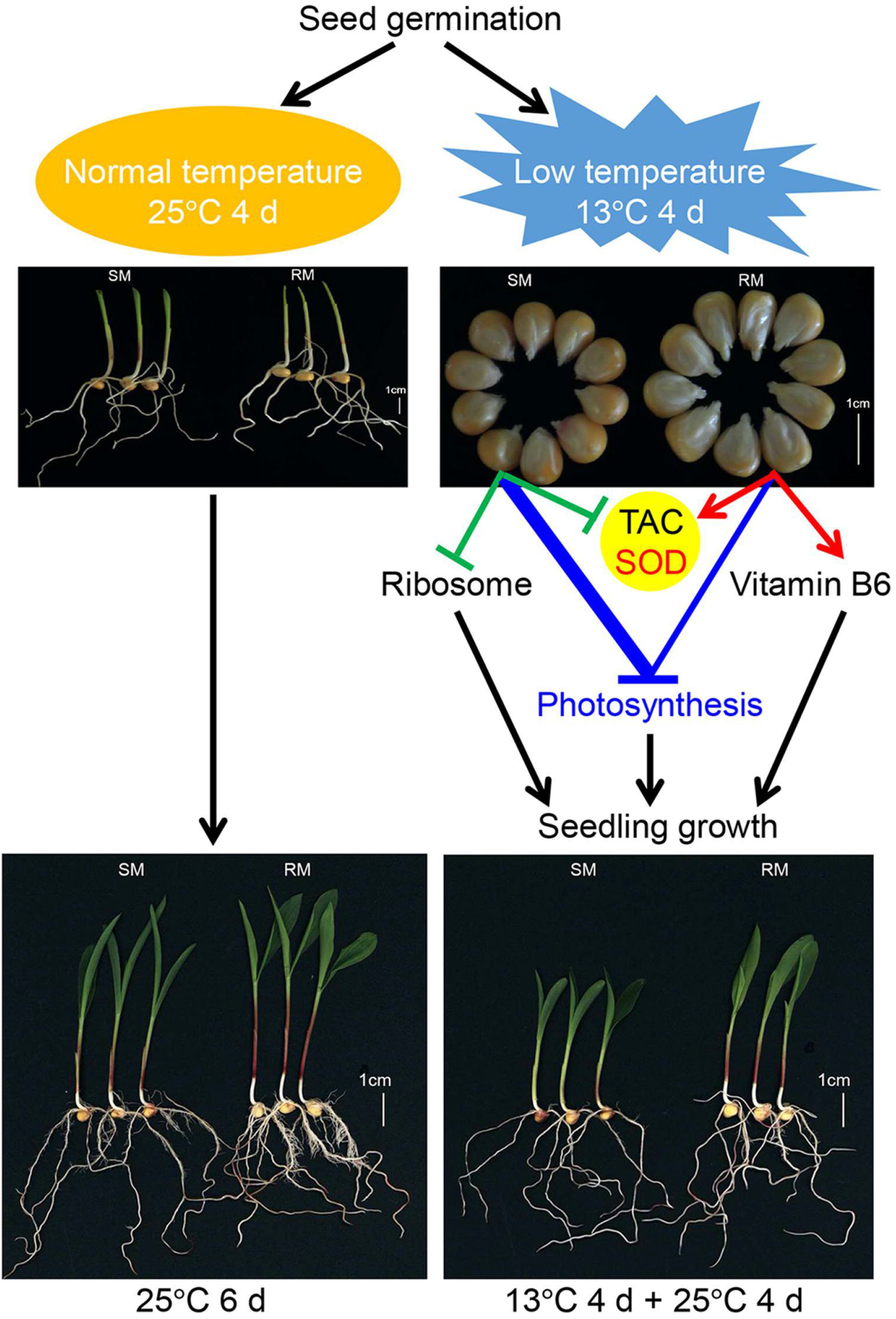
A possible network of maize seed germination under low temperature affecting subsequent seedling growth. The thickened blue line indicates stronger changes induced by low-temperature stress (according to Figure. 5).

## Conclusions

In summary, maize seed germination under low-temperature stress displayed an increase of lipid peroxidation and inhibited subsequent seedling growth under normal temperature. Transcriptome analysis revealed that photosynthesis and antioxidant metabolism related pathways played important roles in seed germination in response to low-temperature stress at the germination stage, and the photosynthetic system of the SM line was more vulnerable to low-temperature stress. Moreover, SOD activity might play a key role in TAC in seed germination under low-temperature stress. Therefore, this study provides new insights into maize seed germination in response to low-temperature stress.

## Materials and methods

### Materials

The low-temperature sensitive maize (SM) B283-1 and low-temperature resistance maize (RM) 04Qun0522-1-1maize inbred lines used in this study were bred in our laboratory. They were grown at the experimental station of Shandong Agricultural University, Shandong, China (approximately 117°90’E longitude and 36°90’N latitude) in June 2019. Considering the influence of seed maturity on germination, seeds were harvested 50 days after pollination.

### Seed germination

Maize seeds were germinated on a sprouting bed according to a previous study with some modifications (Wen et al., 2018). In per germination box, base sand (diameter of 0.05–0.8 mm) consisted of 4 cm height silica sand with 60% saturation moisture content. Randomly selected 30 maize seeds were sowed on the surface of base sand, and then they were covered with 2 cm height silica sand with 60% saturation moisture content. Some germination boxes were placed in a growth chamber at 25°C, 70% relative humidity, illumination conditions of 4000 lux, and 12 h light photoperiod for 4 days. Other germination boxes were placed in a growth chamber for 13°C for 4 days and then 25°C for 2 days. Germination boxes placed at 25°C or13°C for 7 d were used to test germination percentage under normal temperature or low-temperature stress, respectively. A seed was considered as germination when the radicle was similar with seed length and the germ was similar with half of the seed length.

### Measurement of TAC

TAC was measured by 2,2’-azino-bis (3-ethylbenzothiazoline-6-sulfonic acid) diammonium salt (ABTS), ABTS oxidized to form a stable blue-green cationic radical ABTS^+^ (Scalzo et al., 2005) which could be dissolved in water or acid ethanol medium and had the maximum absorption at 734 nm using the ABTS Assay Kit (Comin, Suzhou, China) following to the manufacturer’s protocols. When the tested substance is added to the ABTS^+^ solution, the antioxidant components can react with ABTS^+^ and fade the reaction system. The change of absorbance was detected by 734 nm, and the antioxidant capacity of antioxidants was quantified with Trolox as the control system. Three biological replicates were used for each treatment.

### Measurement of lipid peroxidation

Lipid peroxidation was determined using a commercial kit (Comin, Suzhou, China) as previously described (Checchio et al., 2021). Briefly, Add 0.1 g tissue to the 1 mL extract, take out the 90 μL sample extract, add the reagent-900 μL, and mix the reagent two 300 μL immediately and heated at 95°C in a water bath for 1 h, and then the absorbance of the centrifuged supernatant solution was measured at 532 nm. Results were expressed as nmol per g protein. Three biological replicates were used for each treatment.

### Measurement of sucrose content

The determination principle of sucrose content is that alkali is used to heat the sample together to destroy the reducing sugar (Handel, 1968). Then, sucrose was hydrolyzed to glucose and fructose under acidic conditions, and fructose reacted with resorcinol to form a colored substance with a characteristic absorption peak at 480 nm. Sucrose content was extracted using a sucrose content determination kit (Comin, Suzhou, China) according to the manufacturer’s protocols. Three biological replicates were used for each treatment.

### Measurement of SOD activity

SOD activity was determined by Giannopolitis and Ries (1977). Superoxide anion (O^2−^) is produced by xanthine and xanthine oxidase reaction system, O^2−^ can reduce azoblue tetrazole to form blue methyl, which is absorbed in 560 nm. SOD can scavenge O^2−^, thus inhibiting the formation of methylpyrazine. The darker the blue of the reaction solution, the lower the SOD activity, on the contrary, the higher the SOD activity. SOD was extracted using a SOD determination kit (Comin, Suzhou, China) according to the manufacturer’s protocols. Three biological replicates were used for each treatment.

### Measurement of POD activity

POD catalyzes H_2_O_2_ oxidation of specific substrates and has characteristic light absorption at 470 nm (Zhang et al., 2015). POD was extracted using a POD determination kit (Comin, Suzhou, China) according to the manufacturer’s protocols. Three biological replicates were used for each treatment.

### Measurement of SPAD value

The SPAD value was estimated by the SPAD-502 chlorophyll meter (Konica Minolta Inc., Japan). Three biological replicates were used for each treatment. The SPAD values of two inbred lines growing at 25°C for 6 days or germinating at 13°C for 4 days and transferring to 25°C for 4 days were measured.

### RNA-seq and transcriptome analysis

Seedlings from each replication were pooled and stored at −80°C after quick freezing with liquid nitrogen. Three biological replicates were used for each treatment, resulting in 12 samples. Total RNA was extracted using a biospin plant total RNA extraction kit (Bioflux, Beijing, China) according to the manufacturer’s protocols. Libraries were generated using NEBNext®UltraTM RNA Library Prep Kit for Illumina^®^ (NEB, USA) following manufacturer’s, mRNA library construction, and Illumina sequencing was performed as described previously (Yu et al., 2021).

Index of the reference genome (http://www.maizesequence.org/index.html) was built using Hisat2 v2.0.5 and paired-end clean reads were aligned to the reference genome (B73 v4) using Hisat2 v2.0.5 (Yu et al., 2021). Differential expression analyses of two conditions/groups (three biological replicates per treatment) were performed using the DESeq2 R package (1.16.1). The resulting P-values were adjusted using Benjamini and Hochberg’s approach for controlling the false discovery rate. DEGs with padj <0.05 and |log_2_Fold change|≥1 were considered for further analyses.

To understand the main biological functions of DEGs in maize, we carried out an enrichment analysis of DEGs. ClusterProfiler (3.4.4) software was used to achieve GO enrichment analysis of DEGs (Yu et al., 2012). In this study, the GO term with corrected padj <0.05 was used as the GO term with significant enrichment of DEGs. To understand the main metabolic pathway of DEGs in maize, we carried out the Kyoto Encyclopedia of Genes and Genomes (KEGG) enrichment analysis of DEGs. We use cluster Profiler (3.4.4) Statistical enrichment of DEGs in the KEGG pathway was analyzed by software (Yu et al., 2012). Similarly, the KEGG pathways with corrected padj <0.05 were assigned as significantly enriched pathways.

### qRT-PCR

Primers were designed using the primer-premier software (version 6.0) (Supplemental Table S6). The maize *Actin* gene (*Zm00001d010159*), as an internal control (Li et al., 2019a). Real-time PCR was performed using the ABI StepOne Plus Real-time PCR System (Applied Biosystems, CA, USA) according to the instructions of SYBR®Green Real-time PCR Master Mix (Takara, Dalian, China). Reaction conditions were 95°C for 5 min, followed by 40 cycles of denaturation at 95°C for 10 s, annealing at 58°C for 20 s, and extension at 72°C 30 s. The relative quantitative analysis method of 2^−ΔΔCT^ was used to calculate the relative expression of genes (Livak and Schmittgen, 2001).

### Statistical analysis

Multiple comparisons were performed among different samples using Duncan’s test at the 0.05 significance level. All the tests were performed using SPSS Version 21.0 for Windows (SPSS, Chicago, IL, USA).

## Supplemental Data

Supplemental Table S1. Summary of RNA-seq data.

Supplemental Table S2. List of SM-specific DEGs for ribosome pathway.

Supplemental Table S3. List of RM-specific DEGs that were significantly enriched in KEGG pathways.

Supplemental Table S4. List of common DEGs in SM_LNTvsNT and RM_LNTvsNT mapped to photosynthesis related pathways.

Supplemental Table S5. List of common DEGs in SM_LNTvsNT and RM_LNTvsNT mapped to glutathione metabolism pathway.

Supplemental Table S6. Primers used for qRT-PCR.

## Acknowledgments

This work was supported by the Maize Industry Technology System in Shandong Province (SDAIT-02-02).

## Notes

### Competing Interest Statement

The authors have declared no competing interest.

### Summary of Updates

Figure 8 revised

## References

Bose J, Rodrigo-Moreno A, Shabala S (2014) ROS homeostasis in halophytes in the context of salinity stress tolerance. Journal of Experimental Botany 65: 1241–1257

Cao Q, Li G, Cui Z, Yang F, Jiang X, Diallo L, Kong F (2019) Seed Priming with Melatonin Improves the Seed Germination of Waxy Maize under Chilling Stress via Promoting the Antioxidant System and Starch Metabolism. Scientific Reports 9: 1–12

Checchio MV, de Cássia Alves R, de Oliveira KR, Moro GV, Santos DMM dos, Gratão PL (2021) Enhancement of salt tolerance in corn using Azospirillum brasilense: an approach on antioxidant systems. Journal of Plant Research. doi: 10.1007/s10265-021-01332-1

Cnubben NHP, Rietjens IMCM, Wortelboer H, Zanden J Van, Bladeren PJ Van (2001) The interplay of glutathione-related processes in antioxidant defense. Environmental Toxicology and Pharmacology 10: 141–152

Denslow SA, Rueschhoff EE, Daub ME (2007) Regulation of the Arabidopsis thaliana vitamin B6 biosynthesis genes by abiotic stress. Plant Physiology and Biochemistry 45: 152–161

Drewke C, Leistner E (2001) Biosynthesis of vitamin B6 and structurally related derivatives. Vitamins and Hormones 61: 121–155

Ehrenshaft M, Chung KR, Jenns AE, Daub ME (1999) Functional characterization of SOR1, a gene required for resistance to photosensitizing toxins in the fungus Cercospora nicotianae. Current Genetics 34: 478–485

Ehrenshaft M, Jenns AE, Chung KR, Daub ME (1998) SOR1, a gene required for photosensitizer and singlet oxygen resistance in Cercospora fungi, is highly conserved in divergent organisms. Molecular Cell 1: 603–609

Ensminger I, Busch F, Huner NPA (2006) Photostasis and cold acclimation: Sensing low temperature through photosynthesis. Physiologia Plantarum 126: 28–44

Fenza M Di, Hogg B, Grant J, Barth S (2017) Transcriptomic response of maize primary roots to low temperatures at seedling emergence. PeerJ 2017: 1–17

Garcia-Molina A, Kleine T, Schneider K, Mühlhaus T, Lehmann M, Leister D (2020) Translational Components Contribute to Acclimation Responses to High Light, Heat, and Cold in Arabidopsis. iScience 23: 1–21

Giannopolitis CN and Ries SK (1977) Superoxide dismutases. Plant Physiology 59: 309–314

Grzybowski M, Adamczyk J, Jończyk M, Sobkowiak A, Szczepanik J, Frankiewicz K, Fronk J, Sowiński P (2019) Increased photosensitivity at early growth as a possible mechanism of maize adaptation to cold springs. Journal of Experimental Botany 70: 2887–2904

Gu L, Zhao M, Ge M, Zhu S, Cheng B, Li X (2019) Transcriptome analysis reveals comprehensive responses to cadmium stress in maize inoculated with arbuscular mycorrhizal fungi. Ecotoxicology and Environmental Safety 186: 1–9

Handel E V (1968) Direct microdetermination of sucrose. Analytical Biochemistry 22: 280–283

Hodges DM, Andrews CJ, Johnson DA, Hamilton RI (1997) Antioxidant enzyme and compound responses to chilling stress and their combining abilities in differentially sensitive maize hybrids. Crop Science 37: 857–863

Huang SH, Zhang JY, Wang LH, Huang LQ (2013) Effect of abiotic stress on the abundance of different vitamin B6 vitamers in tobacco plants. Plant Physiology and Biochemistry 66: 63–67

Kupsch C, Ruwe H, Gusewski S, Tillich M, Small I, Schmitz-Linneweber C (2012) Arabidopsis chloroplast RNA binding proteins CP31A and CP29A associate with large transcript pools and confer cold stress tolerance by influencing multiple chloroplast RNA processing steps. Plant Cell 24:4266–4280

Li M, Sui N, Lin L, Yang Z, Zhang Y (2019a) Transcriptomic profiling revealed genes involved in response to cold stress in maize. Functional Plant Biology 46: 830–844

Li Y, Wang X, Ban Q, Zhu X, Jiang C, Wei C, Bennetzen JL (2019b) Comparative transcriptomic analysis reveals gene expression associated with cold adaptation in the tea plant Camellia sinensis. BMC Genomics 20: 1–17

Liu X, Rodermel SR, Yu F (2010) A var2 leaf variegation suppressor locus, SUPPRESSOR OF VARIEGATION3, encodes a putative chloroplast translation elongation factor that is important for chloroplast development in the cold. BMC Plant Biology 10: 1–18

Livak KJ, Schmittgen TD (2001) Analysis of relative gene expression data using real-time quantitative PCR and the 2-ΔΔCT method. Methods 25: 402–408

Ma Y, Dai X, Xu Y, Luo W, Zheng X, Zeng D, Pan Y, Lin X, Liu H, Zhang D, et al (2015) COLD1 confers chilling tolerance in rice. Cell 160: 1209–1221

Nägele T, Heyer AG (2013) Approximating subcellular organisation of carbohydrate metabolism during cold acclimation in different natural accessions of Arabidopsis thaliana. New Phytologist 198:777–787

Nie G □Y, Long SP, Baker NR (1992) The effects of development at sub□optimal growth temperatures on photosynthetic capacity and susceptibility to chilling□dependent photoinhibition in Zea mays. Physiologia Plantarum 85: 554–560

Noctor G, Mhamdi A, Chaouch S, Han Y, Neukermans J, Marquez-Garcia B, Queval G, Foyer CH (2012) Glutathione in plants: An integrated overview. Plant, Cell and Environment 35: 454–484

Ohama N, Sato H, Shinozaki K, Yamaguchi-Shinozaki K (2017) Transcriptional Regulatory Network of Plant Heat Stress Response. Trends in Plant Science 22: 53–65

Pfister AS (2019) Emerging role of the nucleolar stress response in autophagy. Frontiers in Cellular Neuroscience 13: 1–18

Prasad TK (1997) Role of Catalase in lnducing Chilling Tolerance in Pre-emergent Maize Seedlings. Plant Physiology 114: 1369–1376

Pulido P, Zagari N, Manavski N, Gawroński P, Matthes A, Scharff LB, Meurer J, Leister D (2018) CHLOROPLAST RIBOSOME ASSOCIATED supports translation under stress and interacts with the ribosomal 30S subunit. Plant Physiology 177: 1539–1554

Reiter B, Vamvaka E, Marino G, Kleine T, Jahns P, Bolle C, Leister D, Rühle T (2020) The Arabidopsis protein CGL20 is required for plastid 50S ribosome biogenesis. Plant Physiology 182:1222–1238

Romero-Puertas MC, Corpas FJ, Sandalio LM, Leterrier M, Rodríguez-Serrano M, Del Río LA, Palma JM (2006) Glutathione reductase from pea leaves: Response to abiotic stress and characterization of the peroxisomal isozyme. New Phytologist 170: 43–52

Savitch L V., Ivanov AG, Gudynaite-Savitch L, Huner NPA, Simmonds J (2011) Cold stress effects on PSI photochemistry in Zea mays: Differential increase of FQR-dependent cyclic electron flow and functional implications. Plant and Cell Physiology 52: 1042–1054

Scalzo J, Politi A, Pellegrini N, Mezzetti B, Battino M (2005) Plant genotype affects total antioxidant capacity and phenolic contents in fruit. Nutrition 21: 207–213

Schultes NP, Sawers RJH, Brutnell TP, Krueger RW (2000) Maize high chlorophyll fluorescent 60 mutation is caused by an Ac disruption of the gene encoding the chloroplast ribosomal small subunit protein 17. Plant Journal 21: 317–327

Sowiński P, Rudzińska-Langwald A, Adamczyk J, Kubica I, Fronk J (2005) Recovery of maize seedling growth, development and photosynthetic efficiency after initial growth at low temperature. Journal of Plant Physiology 162: 67–80

Wang W, Zheng K, Gong X, Xu J, Huang J, Lin D, Dong Y (2017) The rice TCD11 encoding plastid ribosomal protein S6 is essential for chloroplast development at low temperature. Plant Science 259: 1–11

Wen D, Hou H, Meng A, Meng J, Xie L, Zhang C (2018) Rapid evaluation of seed vigor by the absolute content of protein in seed within the same crop. Scientific Reports 8: 1–8

Xu T, Lee K, Gu L, Kim J Il, Kang H (2013) Functional characterization of a plastid-specific ribosomal protein PSRP2 in Arabidopsis thaliana under abiotic stress conditions. Plant Physiology and Biochemistry 73: 405–411

Yang K, Yang J, Yi J (2018) Nucleolar stress: Hallmarks, sensing mechanism and diseases. Cell Stress 2: 125–140

Yu G, Wang LG, Han Y, He QY (2012) ClusterProfiler: An R package for comparing biological themes among gene clusters. OMICS A Journal of Integrative Biology 16: 284–287

Yu T, Zhang J, Cao J, Cai Q, Li X, Sun Y, Li S, Li Y, Hu G, Cao S, et al (2021) Leaf transcriptomic response mediated by cold stress in two maize inbred lines with contrasting tolerance levels. Genomics 113: 782–794

Zeng R, Li Z, Shi Y, Fu D, Yin P, Cheng J, Jiang C, Yang S (2021) Natural variation in a type-A response regulator confers maize chilling tolerance. Nature Communications 12: 1–13

Zhang H, Zhang J, Xu Q, Wang D, Di H, Huang J, Yang X, Wang Z, Zhang L, Dong L, et al (2020) Identification of candidate tolerance genes to low-temperature during maize germination by GWAS and RNA-seqapproaches. BMC Plant Biology 20: 1–17

Zhang J, Yuan H, Yang Y, Fish T, Lyi SM, Thannhauser TW, Zhang L, Li L (2016) Plastid ribosomal protein S5 is involved in photosynthesis, plant development, and cold stress tolerance in Arabidopsis. Journal of Experimental Botany 67: 2731–2744

Zhang L, Pei Y, Wang H, Jin Z, Liu Z, Qiao Z, Fang H, Zhang Y (2015) Hydrogen sulfide alleviates cadmium-induced cell death through restraining ROS accumulation in roots of Brassica rapa L. ssp. pekinensis. Oxidative Medicine and Cellular Longevity 2015: 1–11

Zhang Q, Bartels D (2018) Molecular responses to dehydration and desiccation in desiccation-tolerant angiosperm plants. Journal of Experimental Botany 69: 3211–3222

Zhao Q, Zhou L, Liu J, Du X, Asad MAU, Huang F, Pan G, Cheng F (2018) Relationship of ROS accumulation and superoxide dismutase isozymes in developing anther with floret fertility of rice under heat stress. Plant Physiology and Biochemistry 122: 90–101

Zheng H, Yang Z, Wang W, Guo S, Li Z, Liu K, Sui N (2020) Transcriptome analysis of maize inbred lines differing in drought tolerance provides novel insights into the molecular mechanisms of drought responses in roots. Plant Physiology and Biochemistry 149: 11–26

Zheng Z, Xu X, Crosley RA, Greenwalt SA, Sun Y, Blakeslee B, Wang L, Ni W, Sopko MS, Yao C, et al (2010) The protein kinase SnRK2.6 mediates the regulation of sucrose metabolism and plant growth in arabidopsis. Plant Physiology 153: 99–113

