## Supplemental Material for "Photosynthesis and antioxidant metabolism modulate the low-temperature resistance of seed germination in maize"

**Supplemental Table S1.** Summary of RNA-seq data.

| Sample | Clean reads | Total mapped | Uniquely mapped | Multiple mapped |
| --- | --- | --- | --- | --- |
| SM_NT-1 | 43490354 | 37836919(87.00%) | 36555093(84.05%) | 1281826(2.95%) |
| SM_NT-2 | 41222002 | 35387748(85.85%) | 34353744(83.34%) | 1034004(2.51%) |
| SM_NT-3 | 43273292 | 38487979(88.94%) | 37179188(85.92%) | 1308791(3.02%) |
| SM_LNT-1 | 50390222 | 45859681(91.01%) | 44568538(88.45%) | 1291143(2.56%) |
| SM_LNT-2 | 44459968 | 40043970(90.07%) | 38926826(87.55%) | 1117144(2.51%) |
| SM_LNT-3 | 41122500 | 37081813(90.17%) | 36054421(87.68%) | 1027392(2.50%) |
| RM_NT-1 | 43804464 | 38580693(88.07%) | 37503118(85.61%) | 1077575(2.46%) |
| RM_NT-2 | 40378292 | 36434609(90.23%) | 35393515(87.65%) | 1041094(2.58%) |
| RM_NT-3 | 44991214 | 40525360(90.07%) | 39320133(87.40%) | 1205227(2.68%) |
| RM_LNT-1 | 44004232 | 38302870(87.04%) | 37106847(84.33%) | 1196023(2.72%) |
| RM_LNT-2 | 43780552 | 39255224(89.66%) | 38090518(87.00%) | 1164706(2.66%) |
| RM_LNT-3 | 39838102 | 35301330(88.61%) | 34317047(86.14%) | 984283(2.47%) |

**Supplemental Table S2.** List of SM-specific DEGs for ribosome pathway.

| Gene ID | Gene description | Log_2_(fold change) |
| --- | --- | --- |
|  |  | SM_LNTvsNT |
| *Zm00001d038835* | 30S ribosomal protein S1, chloroplastic | -1.253 |
| *Zm00001d028153* | 30S ribosomal protein S10 chloroplastic | -2.379 |
| *Zm00001d012353* | 30S ribosomal protein S17 chloroplastic | -2.545 |
| *Zm00001d047186* | 30S ribosomal protein S4, chloroplastic | -1.139 |
| *Zm00001d034808* | 30S ribosomal protein S6 alpha chloroplastic | -2.474 |
| *Zm00001d011816* | 40S ribosomal protein S10-2 | -1.063 |
| *Zm00001d000120* | 40S ribosomal protein S14 | -1.194 |
| *Zm00001d009000* | 40S ribosomal protein S15a-1 | -1.104 |
| *Zm00001d039299* | 40S ribosomal protein S15a-1 | -1.939 |
| *Zm00001d052885* | 40S ribosomal protein S15a-1 | -1.357 |
| *Zm00001d047296* | 40S ribosomal protein S19 | -1.050 |
| *Zm00001d029543* | 40S ribosomal protein S19-3 | -1.233 |
| *Zm00001d045448* | 40S ribosomal protein S20-1 | -1.235 |
| *Zm00001d028426* | 40S ribosomal protein S20-1 | -1.316 |
| *Zm00001d043606* | 40S ribosomal protein S24 | -1.113 |
| *Zm00001d011741* | 40S ribosomal protein S24 | -1.082 |
| *Zm00001d034609* | 40S ribosomal protein S26 | -1.066 |
| *Zm00001d041727* | 40S ribosomal protein S29 | -1.313 |
| *Zm00001d022111* | 40S ribosomal protein S3-1 | 1.023 |
| *Zm00001d037929* | 40S ribosomal protein S5-2 | -1.294 |
| *Zm00001d028786* | 40S ribosomal protein S7 | -1.065 |
| *Zm00001d047707* | 40S ribosomal protein S7 | -1.049 |
| *Zm00001d053633* | 40S ribosomal protein S8 | -1.262 |
| *Zm00001d038084* | 50S ribosomal protein L1 chloroplastic | -2.630 |
| *Zm00001d027421* | 50S ribosomal protein L11 chloroplastic | -2.871 |
| *Zm00001d022197* | 50S ribosomal protein L12-2 | 1.411 |
| *Zm00001d041322* | 50S ribosomal protein L14 | -1.252 |
| *Zm00001d012998* | 50S ribosomal protein L17 chloroplastic | -3.060 |
| *Zm00001d010803* | 50S ribosomal protein L20 | -1.041 |
| *Zm00001d053377* | 50S ribosomal protein L21 chloroplastic | -2.060 |
| *Zm00001d044130* | 50S ribosomal protein L31 | -2.751 |
| *Zm00001d047462* | 50S ribosomal protein L6 chloroplastic | -1.959 |
| *Zm00001d013730* | 5S rRNA binding protein | -1.065 |
| *Zm00001d009652* | 60S acidic ribosomal protein P2-5 | -1.203 |
| *Zm00001d026578* | 60S acidic ribosomal protein P2A | -1.079 |
| *Zm00001d046317* | 60S ribosomal protein L11-1 | -1.087 |
| *Zm00001d010016* | 60S ribosomal protein L11-1 | -1.035 |
| *Zm00001d017719* | 60S ribosomal protein L12-3 | -1.886 |
| *Zm00001d011992* | 60S ribosomal protein L13 | -1.050 |
| *Zm00001d042308* | 60S ribosomal protein L13 | -1.003 |
| *Zm00001d013252* | 60S ribosomal protein L13a-1 | -1.106 |
| *Zm00001d006287* | 60S ribosomal protein L14-1 | -1.367 |
| *Zm00001d025857* | 60S ribosomal protein L14-1 | -1.341 |
| *Zm00001d011534* | 60S ribosomal protein L18a | -1.158 |
| *Zm00001d048382* | 60S ribosomal protein L21-1 | -1.371 |
| *Zm00001d002462* | 60S ribosomal protein L22-2 | 1.460 |
| *Zm00001d007154* | 60S ribosomal protein L22-2 | -1.174 |
| *Zm00001d048368* | 60S ribosomal protein L23 | -1.016 |
| *Zm00001d037489* | 60S ribosomal protein L23a-1 | -1.669 |
| *Zm00001d013758* | 60S ribosomal protein L24 | -1.785 |
| *Zm00001d010631* | 60S ribosomal protein L24 | -1.069 |
| *Zm00001d015375* | 60S ribosomal protein L27a-3 | -1.227 |
| *Zm00001d009417* | 60S ribosomal protein L28-1 | -1.283 |
| *Zm00001d040617* | 60S ribosomal protein L29 | -1.158 |
| *Zm00001d038535* | 60S ribosomal protein L30-2 | -1.106 |
| *Zm00001d017850* | 60S ribosomal protein L31 | -1.317 |
| *Zm00001d031429* | 60S ribosomal protein L31 | -2.076 |
| *Zm00001d051662* | 60S ribosomal protein L31 | -1.603 |
| *Zm00001d045889* | 60S ribosomal protein L31 | -1.068 |
| *Zm00001d049790* | 60S ribosomal protein L32 | -1.080 |
| *Zm00001d021018* | 60S ribosomal protein L32-1 | -1.032 |
| *Zm00001d024571* | 60S ribosomal protein L34 | -1.140 |
| *Zm00001d020450* | 60S ribosomal protein L34-3 | -1.527 |
| *Zm00001d005642* | 60S ribosomal protein L35a-2 | -1.651 |
| *Zm00001d010528* | 60S ribosomal protein L36 | -1.734 |
| *Zm00001d038374* | 60S ribosomal protein L36 | -1.591 |
| *Zm00001d006510* | 60S ribosomal protein L44 | -1.633 |
| *Zm00001d019562* | 60S ribosomal protein L44 | -1.115 |
| *Zm00001d027512* | 60S ribosomal protein L5-1 | -1.033 |
| *Zm00001d048181* | Chloroplast 30S ribosomal protein S10 | -2.441 |
| *Zm00001d016200* | N/A | -1.326 |
| *Zm00001d009117* | N/A | -1.074 |
| *Zm00001d017449* | Ribosomal protein L19 | -3.105 |
| *Zm00001d044287* | Ribosomal protein L19 | -1.350 |
| *Zm00001d045069* | Ribosomal protein L19 | -1.296 |
| *Zm00001d014969* | Ribosomal protein L37 | -1.242 |
| *Zm00001d016276* | Ribosomal protein S24 | -1.495 |
| *Zm00001d049666* | Ribosomal protein13 | -1.201 |

**Supplemental Table S3**. List of RM-specific DEGs that were significantly enriched in KEGG pathways.

| Gene ID | Gene description | Log_2_(fold change) |
| --- | --- | --- |
|  |  | RM_LNTvsNT |
| **Phenylpropanoid biosynthesis** | | |
| *Zm00001d002901* | Peroxidase 12 | -5.024 |
| *Zm00001d003707* | Peroxidase 72 | -2.329 |
| *Zm00001d004443* | Probable cinnamyl alcohol dehydrogenase 9 | -2.091 |
| *Zm00001d007952* | Peroxidase 73 | 5.200 |
| *Zm00001d008435* | Beta-glucosidase 44 | 2.965 |
| *Zm00001d015459* | 4-coumarate--CoA ligase 1 | -1.601 |
| *Zm00001d017274* | Phenylalanine ammonia-lyase | -1.823 |
| *Zm00001d017275* | phenylalanine ammonia lyase9 | -1.494 |
| *Zm00001d017279* | phenylalanine ammonia lyase7 | -2.376 |
| *Zm00001d018620* | Peroxidase 1 | -2.496 |
| *Zm00001d021533* | Peroxidase 7 | -4.569 |
| *Zm00001d022278* | 1-Cys peroxiredoxin PER1 | 5.619 |
| *Zm00001d022453* | Peroxidase 52 | -5.856 |
| *Zm00001d037547* | guaiacol peroxidase3 | -2.840 |
| *Zm00001d039837* | Spermidine hydroxycinnamoyl transferase | -1.555 |
| *Zm00001d039840* | hydroxycinnamoyltransferase7 | -5.176 |
| *Zm00001d040364* | Peroxidase 72 | -2.536 |
| *Zm00001d044339* | aldehyde dehydrogenase3 | -1.217 |
| *Zm00001d044340* | aldehyde dehydrogenase5 | -2.985 |
| *Zm00001d009373* | Peroxidase | -1.438 |
| *Zm00001d014467* | Peroxidase | -2.001 |
| *Zm00001d016471* | Trans-cinnamate 4-monooxygenase | 1.394 |
| *Zm00001d020400* | Cinnamyl alcohol dehydrogenase1 | -3.025 |
| *Zm00001d025402* | Peroxidase | -2.535 |
| *Zm00001d027710* | Peroxidase | -3.488 |
| *Zm00001d032148* | Hydroxycinnamoyltransferase11 | -1.857 |
| *Zm00001d034015* | Exoglucanase1 | -1.298 |
| *Zm00001d039833* | Putrescine hydroxycinnamoyltransferase 1 | -5.731 |
| *Zm00001d040705* | Peroxidase | -1.406 |
| *Zm00001d045043* | Putative cinnamyl alcohol dehydrogenase 1 | 1.499 |
| *Zm00001d047514* | Peroxidase | -3.233 |
| *Zm00001d051166* | Phenylalanine ammonia-lyase | -1.349 |
| *Zm00001d051529* | N/A | -1.155 |
| *Zm00001d006937* | Peroxidase | -4.469 |
| **Phenylalanine metabolism** | |  |
| *Zm00001d006638* | Amine oxidase | -1.343 |
| *Zm00001d016198* | glutamate-oxaloacetic transaminase3 | 1.294 |
| *Zm00001d017274* | Phenylalanine ammonia-lyase | -1.823 |
| *Zm00001d017275* | phenylalanine ammonia lyase9 | -1.494 |
| *Zm00001d017279* | phenylalanine ammonia lyase7 | -2.376 |
| *Zm00001d025103* | Amine oxidase | 2.976 |
| *Zm00001d043382* | glutamate-oxaloacetate transaminase1 | 2.519 |
| *Zm00001d051166* | Phenylalanine ammonia-lyase | -1.349 |
| **Ubiquinone and other terpenoid-quinone biosynthesis** | | |
| *Zm00001d005251* | NAD(P)H dehydrogenase (quinone) | -2.112 |
| *Zm00001d015459* | 4-coumarate--CoA ligase 1 | -1.601 |
| *Zm00001d017746* | Vitamin E synthesis4 | 2.332 |
| *Zm00001d037542* | NAD(P)H dehydrogenase (quinone) | -3.528 |
| *Zm00001d045322* | Ubiquinone biosynthesis O-methyltransferase, mitochondrial | 1.402 |
| *Zm00001d016471* | Trans-cinnamate 4-monooxygenase | 1.394 |
| *Zm00001d024771* | 1,4-dihydroxy-2-naphthoyl-CoA thioesterase 1 | 1.568 |
| *Zm00001d051529* | N/A | -1.155 |
| **Tropane, piperidine and pyridine alkaloid biosynthesis** | | |
| *Zm00001d043382* | glutamate-oxaloacetate transaminase1 | 2.519 |
| *Zm00001d016198* | glutamate-oxaloacetic transaminase3 | 1.294 |
| *Zm00001d006638* | Amine oxidase | -1.343 |
| *Zm00001d007687* | Tropinone reductase-like protein | 1.189 |
| *Zm00001d025103* | Amine oxidase | 2.976 |

**Supplemental Table S4**. List of common DEGs in SM_LNTvsNT and RM_LNTvsNT mapped to photosynthesis related pathways.

| Gene ID | Gene description | Log_2_(fold change) | | |
| --- | --- | --- | --- | --- |
|  |  | SM_LNTvsNT | | RM_LNTvsNT |
| **Photosynthesis** | | | | |
| *Zm00001d003767* | Photosystem I subunit O | -6.244 | -1.092 | |
| *Zm00001d005446* | Photosystem I reaction center subunit IV A | -4.266 | -1.344 | |
| *Zm00001d005996* | Photosystem I reaction center subunit V | -4.667 | -2.369 | |
| *Zm00001d007857* | Oxygen-evolving enhancer protein 2-1 chloroplastic | -3.473 | -1.330 | |
| *Zm00001d012293* | Probable ferredoxin-4 chloroplastic | 6.685 | -4.043 | |
| *Zm00001d013146* | Photosystem I reaction center subunit III chloroplastic | -3.349 | -1.567 | |
| *Zm00001d016134* | Iron-sulfur protein1 | -3.561 | -1.501 | |
| *Zm00001d018069* | ATP synthase chloroplast subunit1 | -3.664 | -1.398 | |
| *Zm00001d018779* | Oxygen-evolving enhancer protein 2-1 chloroplastic | -3.864 | -1.099 | |
| *Zm00001d018797* | photosystem I reaction center6 | -3.932 | -1.743 | |
| *Zm00001d020877* | Photosystem I reaction center subunit V chloroplastic | -3.518 | -2.628 | |
| *Zm00001d021620* | ATP synthase chloroplast subunit2 | -3.271 | -1.017 | |
| *Zm00001d021703* | Oxygen evolving complex2 | -3.489 | -1.640 | |
| *Zm00001d023713* | Photosystem I reaction center subunit N | -3.034 | -3.770 | |
| *Zm00001d029049* | Photosystem II repair protein PSB27-H1 chloroplastic | -3.343 | -1.536 | |
| *Zm00001d034283* | Photosystem I reaction center subunit III | -3.644 | -1.710 | |
| *Zm00001d034543* | Photosystem I reaction center subunit II-1 chloroplastic | -3.688 | -1.650 | |
| *Zm00001d035001* | Ferredoxin1 | -1.764 | -3.464 | |
| *Zm00001d035002* | Ferredoxin5 | -1.770 | -2.688 | |
| *Zm00001d035003* | Ferredoxin2 | -5.217 | -2.453 | |
| *Zm00001d035135* | Photosystem II core complex protein psbY | -1.235 | -3.075 | |
| *Zm00001d038984* | photosystem I H subunit1 | -4.414 | -1.720 | |
| *Zm00001d039687* | Photosystem I reaction center subunit XI chloroplastic | 5.088 | -8.928 | |
| *Zm00001d042049* | Ferredoxin | -4.020 | -1.506 | |
| *Zm00001d042178* | Photosystem II reaction center psb28 protein | -1.503 | -3.833 | |
| *Zm00001d042697* | photosystem II subunit PsbS1 | -3.673 | -3.597 | |
| *Zm00001d049732* | Ferredoxin-1 | -6.634 | -2.444 | |
| *Zm00001d052242* | ATP synthase subunit delta chloroplastic | -3.486 | -1.129 | |
| *Zm00001d053432* | Iron-sulfur protein2 | -3.053 | -2.180 | |
| *Zm00001d011833* | Ferredoxin--NADP reductase leaf isozyme 1 chloroplastic | -3.130 | -1.444 | |
| *Zm00001d025874* | Uncharacterized protein | 6.722 | -2.913 | |
| *Zm00001d050889* | Oxygen evolving enhancer protein 3 | -7.237 | -3.207 | |
| **Photosynthesis - antenna proteins** | | | | |
| *Zm00001d006587* | Light harvesting complex photosystem II subunit 6 | -3.407 | -1.529 | |
| *Zm00001d001857* | Photosystem I chlorophyll a/b-binding protein 6 chloroplastic | -3.069 | -2.615 | |
| *Zm00001d005814* | Chlorophyll a-b binding protein CP29.1 chloroplastic | -3.123 | -3.439 | |
| *Zm00001d006663* | Light harvesting complex A1 | -4.533 | -2.178 | |
| *Zm00001d007267* | Light harvesting chlorophyll a/b binding protein5 | -3.507 | -1.560 | |
| *Zm00001d009589* | Chlorophyll a-b binding protein chloroplastic | -3.533 | -2.433 | |
| *Zm00001d011285* | Photosystem II light harvesting complex gene B1B2 | -5.026 | -2.871 | |
| *Zm00001d015385* | Chlorophyll a-b binding protein 6 chloroplastic | -3.840 | -1.944 | |
| *Zm00001d018157* | Light harvesting complex a/b protein4 | -4.625 | -1.177 | |
| *Zm00001d021435* | Chlorophyll a-b binding protein chloroplastic | -2.429 | -1.534 | |
| *Zm00001d021763* | Photosystem II subunit29 | -3.507 | -1.313 | |
| *Zm00001d021906* | Chlorophyll a-b binding protein | -5.769 | -1.223 | |
| *Zm00001d026599* | Light harvesting chlorophyll a/b binding protein6 | -3.554 | -2.239 | |
| *Zm00001d032197* | Chlorophyll a-b binding protein 4 chloroplastic | -3.148 | -2.773 | |
| *Zm00001d033136* | Light harvesting chlorophyll binding protein9 | -3.153 | -2.231 | |
| *Zm00001d039040* | Light harvesting complex mesophyll7 | -2.728 | -2.203 | |
| *Zm00001d044396* | Chlorophyll a-b binding protein chloroplastic | -5.063 | -4.064 | |
| *Zm00001d044402* | Chlorophyll a-b binding protein 2 | -4.764 | -2.301 | |
| *Zm00001d046786* | Photosystem I chlorophyll a/b-binding protein 3-1 chloroplastic | -3.587 | -2.073 | |
| *Zm00001d048998* | Chlorophyll a-b binding protein CP26 chloroplastic | -4.047 | -1.413 | |
| *Zm00001d050403* | Chlorophyll a-b binding protein 4 | -4.650 | -1.903 | |
| **Porphyrin and chlorophyll metabolism** | | | | |
| *Zm00001d026405* | Glutamyl-tRNA reductase | -3.395 | -1.223 | |
| *Zm00001d002951* | Protein STAY-GREEN LIKE chloroplastic | -3.574 | -1.311 | |
| *Zm00001d008230* | Magnesium-protoporphyrin IX monomethyl ester | -3.194 | -1.087 | |
| *Zm00001d011819* | Chlorophyllide a oxygenase chloroplastic | -2.699 | -1.595 | |
| *Zm00001d018034* | Geranylgeranyl hydrogenase1 | -1.725 | - | |
| *Zm00001d029150* | Divinyl reductase1 | -2.526 | -1.404 | |
| *Zm00001d023536* | Magnesium-chelatase subunit ChlI-1 chloroplastic | -2.703 | -1.255 | |
| *Zm00001d003935* | Uncharacterized protein | 1.282 | -1.331 | |
| *Zm00001d029074* | Uroporphyrinogen decarboxylase 2 chloroplastic | -2.163 | -1.439 | |
| *Zm00001d013013* | Magnesium-chelatase subunit ChlD chloroplastic | -1.474 | -1.297 | |

**Supplemental Table S5**. List of common DEGs in SM_LNTvsNT and RM_LNTvsNT mapped to glutathione metabolism pathway.

| Gene ID | Gene description | Log_2_(fold change) | | |
| --- | --- | --- | --- | --- |
|  |  | SM_LNTvsNT | | RM_LNTvsNT |
| *Zm00001d002704* | Glutathione peroxidase | | -1.454 | 1.982 |
| *Zm00001d010870* | Glutathione S-transferase F13 | | 9.748 | -9.940 |
| *Zm00001d018220* | Glutathione transferase24 | | -2.429 | 2.579 |
| *Zm00001d020780* | Glutathione transferase23 | | 3.107 | -2.511 |
| *Zm00001d021470* | Glutathione transferase16 | | 2.077 | -2.540 |
| *Zm00001d023582* | L-ascorbate peroxidase S chloroplastic/mitochondrial | | 1.184 | -1.122 |
| *Zm00001d026154* | Glutathione peroxidase | | -1.693 | 3.058 |
| *Zm00001d027541* | Glutathione S-transferase 6 | | -1.590 | -1.623 |
| *Zm00001d027557* | Glutathione transferase31 | | -5.217 | 5.655 |
| *Zm00001d029706* | Glutathione transferase39 | | -3.245 | 3.509 |
| *Zm00001d029707* | Glutathione S-transferase GST 38 | | 1.951 | -1.677 |
| *Zm00001d029708* | Glutathione transferase30 | | -3.086 | 1.997 |
| *Zm00001d042102* | Glutathione transferase28 | | -3.138 | 2.363 |
| *Zm00001d043344* | Glutathione transferase8 | | -1.522 | 1.387 |
| *Zm00001d044021* | Isocitrate dehydrogenase | | -1.122 | 1.500 |
| *Zm00001d048354* | Glutathione S-transferase F9 | | -2.371 | -1.516 |
| *Zm00001d048559* | Glutathione transferase35 | | 4.831 | -3.238 |

**Supplemental Table S6**. Primers used for qRT-PCR.

| Primer | Sequence (5' to 3') |
| --- | --- |
| *Zm00001d010159Actin-F* | GCTACGAGATGCCTGATGGTC |
| *Zm00001d010159Actin-R* | CCCCCACTGAGGACAACG |
| *Zm00001d002611-F* | ACCGATTCCCATCCAACCAACC |
| *Zm00001d002611-R* | CCGAGTCCCGCTGCTTATCATC |
| *Zm00001d025103-F* | GACAACTCGTTCGTTCGC |
| *Zm00001d025103-R* | TCTGATTGTTTGTGAAAGCACC |
| *Zm00001d027557-F* | GGTGAAGGCGGTGGAGAAGATC |
| *Zm00001d027557-R* | ACTTGGCGGGAGCATTGATAGG |
| *Zm00001d027422-F* | CGACAAGAAGGCTGGCTACTCT |
| *Zm00001d027422-R* | CGGAGGACCAAGATCACGGATA |
| *Zm00001d021906-F* | TCGTCGTCGTTGTTCTCCTCCT |
| *Zm00001d021906-R* | CCCAGGGATCAAAGCCGAAGTC |
| *Zm00001d046170-F* | ATCACTGACGACGACAAG |
| *Zm00001d046170-R* | ATCCAAGAAGAGAACCGAAT |
